# Sleep-learning Impairs Subsequent Awake-learning

**DOI:** 10.1101/2020.07.16.206482

**Authors:** Simon Ruch, Marc Alain Züst, Katharina Henke

## Abstract

Although we can learn new information while asleep, we cannot consciously remember the sleep-formed memories because learning occurred in an unconscious state. Here, we ask whether sleep-learning expedites the subsequent awake-learning of the same information. To answer this question, we reanalyzed data (Züst et al., 2019, Curr Biol) from napping participants, who learned new semantic associations between fake foreign-words and translation-words (*guga*-ship) while in slow-wave sleep. They retrieved sleep-formed associations unconsciously on an implicit memory test following awakening. Then, participants took five runs of paired-associative learning to probe carry-over effects of sleep-learning on awake-learning. Surprisingly, sleep-learning diminished awake-learning when participants learned semantic associations that were congruent to sleep-learned associations (*guga*-boat). Yet, learning associations that conflicted with sleep-learned associations (*guga*-coin) was unimpaired relative to learning new associations (*resun*-table; baseline). We speculate that the impeded wake-learning originated in a deficient synaptic downscaling and resulting synaptic saturation in neurons that were activated during both sleep-learning and awake-learning.

## 1 Introduction

Growing evidence suggests that we can unconsciously process and store new information during deep sleep and that this information transfers into wakefulness (Andrillon et al., 2017; Arzi et al., 2012, 2014; Ruch et al., 2014; Ruch & Henke, 2020; Züst et al., 2019). Following awakening, we cannot consciously remember our sleep-formed memories because we formed those memories in the unconscious state of deep sleep. Sleep-formed memories merely exert implicit, indirect effects on our awake behavior (Ruch & Henke, 2020). This may limit the usefulness of sleep-learning for conscious information processing.

Here, we explore whether sleep-learning might expedite the subsequent awake-learning of the same information that had previously been sleep-played. We figured that a first bout of carving a new memory trace during sleep might provide a basis, which ensuing awake-learning can build on. To test this hypothesis, we analyzed previously unreported data acquired in a project on vocabulary learning during slow-wave sleep (SWS) (Züst et al., 2019). These authors observed that new vocabulary was encoded and stored for the long-term during SWS, if the formation of new semantic associations between fake foreign-words and translation-words coincided with peaks of sleep slow-waves (i.e., the positive half waves of slow-waves identified in the electroencephalogram; Fig. 1A). Because the formation of a semantic association cannot take place before the second word is being played, sleep-learning can only be effective when the second word of a pair - which is the time when associative learning occurs - coincides with a brief period of increased cortical excitability and plasticity, i.e., a slow-wave peak. Neuronal network properties resemble those of the waking state during a slow-wave peak (Cox et al., 2014; Destexhe et al., 2007; Schabus et al., 2012; Sirota & Buzsáki, 2005). Peak-associated paired-associative learning yielded a significant associative retrieval performance on the implicit memory test given after awakening (Fig. 1C). Interestingly, sleep-learning, although unconscious, appeared to have recruited the episodic memory system. This was suggested by neuroimaging data collected during implicit memory testing that showed activity increases in the hippocampus and left hemisphere language areas during correctly versus incorrectly responded retrieval trials (Züst et al., 2019). Hence, vocabulary learning during sleep might recruit the same hippocampal-neocortical network that is also involved in vocabulary learning during wakefulness (Breitenstein et al., 2005).

**Figure 1.**
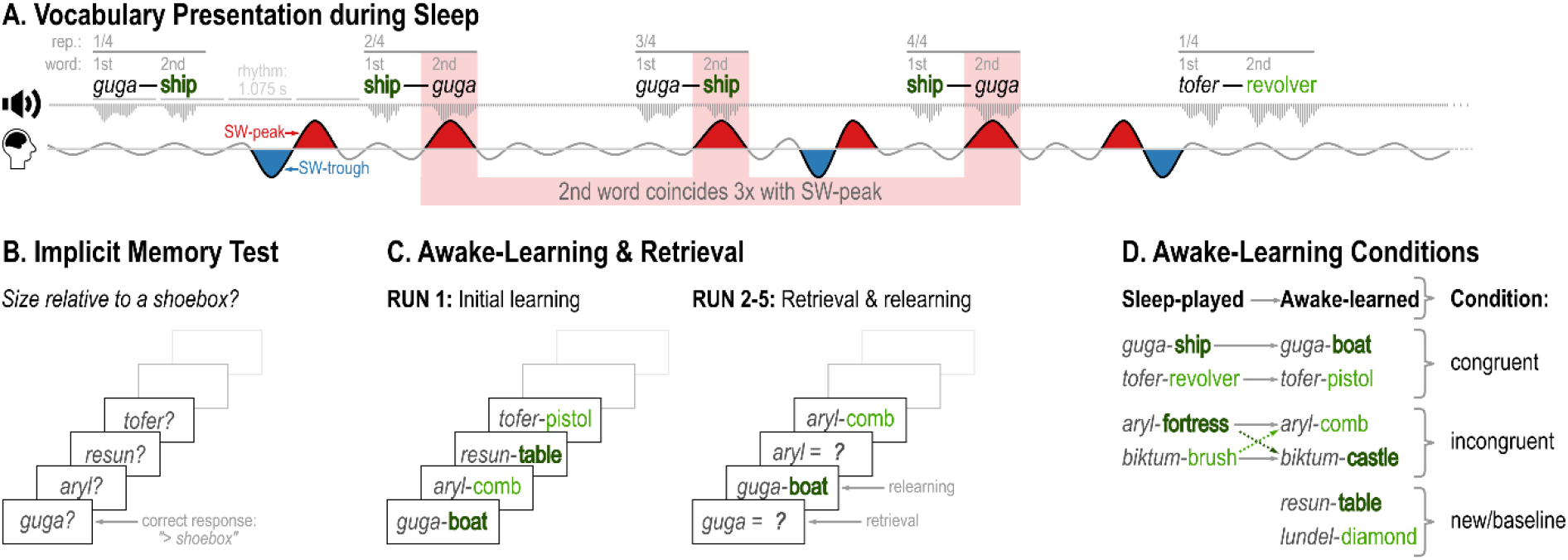
Design. A) Pairs of fake foreign-words and translations-words were presented at a steady rhythm (word onset interval: 1.075 ms, pair onset interval: 4.3 s) during the slow-wave sleep contained in a nap. We played each word-pair four times in sequence, swapping the position of words. Then, we counted the number of times the second word of a pair coincided with a slow-wave peak (red) across the four repetitions of a pair (never, once, twice, three times, or four times). B) Implicit memory test applied following awaking to probe the unconscious reactivation of sleep-learned associations. Participants guessed whether foreign-words designated objects that are larger or smaller than a shoebox. Responses were scored correct if they matched the category (“large” (dark green) or “small” (light green)) of the sleep-played translation-words. C) Awake-learning of new word-pairs (baseline condition) and of previously sleep-played foreign-words combined with translations that are either congruent or incongruent to these words’ sleep-played translations. Run 1: initial learning run; Runs 2-5: cued retrieval with feedback for relearning. D) Overview of awake-learning conditions: congruent, incongruent, and baseline.

If the design of this vocabulary learning task invokes the episodic memory system, then conscious episodic learning in the waking state should profit from previous sleep-learning because both sleep- and awake-learning take place within the same memory system and potentially even within the same neural assemblies. We tested for such a dual-learning benefit by giving participants five consecutive runs of conscious paired-associate vocabulary learning following their sleep-learning and implicit memory testing (Fig. 1B). Multiple runs of awake-learning (and retrieval) allowed delineating the steepness of awake-learning curves. We implemented three conditions of awake-learning (Fig. 1C): In the *semantically congruent condition,* we re-presented sleep-played foreign-words combined with synonyms to their sleep-played translation-words. We used synonyms rather than the sleep-played translation-words to avoid sole perceptual priming and increase chances of reactivating sleep-formed associations in episodic memory by demanding a flexible semantic conversion (*guga*-ship to *guga*-boat). Despite this semantic conversion, memory traces still build on each other because repeated learning in episodic memory relies on the meaning of items and strengthens memories when the material is congruent between learning runs (Höltje et al., 2019; Staresina et al., 2009; van Kesteren et al., 2010, 2013). In the *semantically incongruent condition*, we expected the flattest learning curve because semantically conflicting translation-words were combined with sleep- and awake-presented foreign-words, which creates memory interference and lowers retrieval probability. In this condition, we combined the awake-presented foreign-words with synonyms to other sleep-played translation-words. In the *baseline condition,* we presented new pairs of foreign-words and translation-words to obtain an awake-learning curve that was uninfluenced by previous sleep-learning.

## 2 Materials and Methods

### 2.1 Participants

We report the data of those 26 participants (age 19-32, mean ± SD = 22.96 ± 3.41; 18 (69%) female), who were included in the behavioral part of the original study (Züst et al., 2019). Only those participants underwent the awake-learning procedure. The other group of 15 participants underwent neuroimaging during the implicit memory test without subsequent awake-learning.

The included participants were presented with vocabulary during sleep, took an implicit memory test following awakening, and engaged in conscious paired-associative learning over five runs. This study was approved by the local ethics committee “Kantonale Ethikkommission Bern”.

### 2.2 General procedure

Participants arrived at the sleep laboratory at noon, after a night of partial sleep restriction. They gave written informed consent, were outfitted with EEG electrodes and with in-ear headphones, and then went to sleep for an afternoon nap. Throughout the entire nap, Brownian noise was presented via headphones to reduce the salience of vocabulary presentations. Once the EEG showed clearly visible slow-wave activity (SWS or N2 sleep at the transition to SWS), we initiated the acoustic presentation of word-pairs during sleep. Half of the sleep-played translation-words designated objects that are larger than a shoebox (e.g. ship) and half objects that are smaller than a shoebox (e.g. brush). Each word-pair was repeated four times in sequence with swapped positions (*guga*-ship, ship-*guga*, *guga*-ship, ship-*guga*) to induce a flexible representation of the semantic word-word associations in episodic memory. Words were presented at a steady rhythm (onset interval between words of a pair: 1.075 ms; onset interval between pairs: 4.3 s).

Following the sleep-presentation of the entire set of word-pairs, we woke up participants and gave them time to recover from sleep inertia. Then, participants took an implicit memory test. This test required them to guess whether previously sleep-played foreign-words and new foreign-words designated objects that are larger or smaller than a shoebox. Responses were scored correct if they matched the category (large/small) of the sleep-played translation-words. Responses to new foreign-words were not scored.

Next, we asked participants to engage in five runs of paired-associative vocabulary learning. Half of the sleep-played foreign-words were combined with synonyms to their sleep-played translations (congruent condition) and half were paired with synonyms to other sleep-played translation-words that came from the opposite semantic category (large/small) in order to induce a conflict (incongruent condition; Fig. 1C). New foreign-words that had not been played during sleep were presented with new translation-words to probe uninfluenced paired-associative learning (baseline condition). During the first of five learning runs, all word-pairs were presented for silent learning. In runs 2 to 5, only the foreign-words were presented for participants to cued-recall and produce the translation words. Irrespective of the correctness of the response, the correct translation word was then presented for further paired-associative learning. Participants’ verbal responses were recorded for later analysis. The experimenter proceeded to the feedback trial when a participant had produced a response or – if no response was given – after 5s. The order of word-pairs was re-randomized on each run. Words were presented simultaneously visually on screen and acoustically via loudspeakers. At the end of the experiment, we asked participants whether they had heard anything while asleep. No participant reported having heard words or voices while asleep. Hence, sleep-played words were not perceived with conscious awareness.

### 2.3 Stimulus material

We generated two-syllabic pseudowords (e.g. *guga)* that we used as foreign-words of a fake language. Using a fake foreign language guaranteed that semantic paired-associative learning in this study was uninfluenced by multilingualism. We compiled three lists of 24 foreign-words each. Each list contained 24 synonym pairs that were used as translation-words for sleep-learning and ensuing awake-learning. Half of the synonym-pairs in each list denoted large objects (ship-boat), and half small objects (revolver-pistol). Lists were matched regarding word length, pronounceability, perceived concreteness of foreign-words, and lexical frequency of translation-words. Lists were randomly assigned to the awake-learning conditions (congruent, incongruent, baseline) and counterbalanced over conditions. Within each list, pairs of foreign-words and translation-words were generated randomly for each participant. Translation-words selected for presentation during sleep (e.g. “ship” of “ship-boat”) were never used as synonyms during awake-learning.

### 2.4 Stimulus inclusion and exclusion

Word-pairs intended for sleep-learning were excluded from the data analysis if they were played in a sleep stage other than slow-wave rich N2 sleep or SWS, or during an arousal period.

### 2.5 Electroencephalography

Züst et al. (2019) had found that chances increased that sleep-learning yielded long-term memory traces, if sleep slow-waves peaked between −275 and 25 ms of the onset of the second word in a pair. Because every word-pair was played four times in sequence, the second word could coincide once, twice, three times, four times, or never with a peak. Because awake-learning is bound to be influenced by the robustness of previous sleep-learning, we analyzed the steepness of the awake-learning curve by accounting for the number of slow-wave peaks that had coincided with the play of the second word of a pair. Here is how we determined whether the second word of a pair coincided with a slow-wave peak: We first referenced the raw EEG signal (64 channels, 10-20 montage, 500 Hz sampling rate) to the global average and then extracted the average signal over frontal electrodes (F1, F2, Fz, FC1, FC2, FCz). Then, we low-pass filtered the resulting signal at 4 Hz and computed a time-frequency decomposition (Morlet wavelet transformations, two cycles in length) to extract the instantaneous phase of 0.8 Hz oscillations at 20 ms intervals. The second word of a pair was determined to coincide with a slow-wave peak, if the phase of 0.8 Hz oscillations was within ± 2.5% (± 9°) of the peak phase in the time interval from 275 ms before to 25 ms following the onset of the second word. We determined the number of trials, in which the second word of a pair hit a slow-wave peak several times (2-4 times) and rarely (0/1 times) to contrast these two categories with respect to the later awake-learning success.

### 2.6 Statistical analysis

We performed logistic regressions at the single trial level to find out how retrieval success on the implicit memory test and retrieval success on the runs given for awake-learning would disperse with respect to the number of slow-wave peak-associated word presentations (0/1 times versus 2-4 times) and awake-learning condition (congruent/incongruent/baseline). The binary outcomes were guessing accuracy (correct/incorrect) on the implicit memory test and recall success (yes/no) on each awake-learning run. We modeled random intercepts for participants and foreign-words (mixed effects regressions) when considering the awake-learning runs because both variables accounted significantly for variance in the awake-learning task. Analyses were performed in R (v4.0.2; R Core Team, 2020). We fitted binomial generalized linear models (“glm” function, R-package “stats”) or mixed models (“glmer” function, R-package “lme4”, v1.1-23; Bates et al., 2015) with a logit link.

We fitted full models that included all relevant predictors and all possible interaction terms. To assess the significance of each term, we performed type-II analyses of variance and reported Wald X_W_^2^ statistics (“Anova” function, R-package “car”, v3.0-9; Fox & Weisberg, 2018). We further computed Bayes factors for each effect by analyzing how the fit of a Bayesian binary logistic (mixed) regression model improved when the effect of interest was added. Bayesian models were fitted with the R-package “brms” (v2.14.0) (Bürkner, 2017, 2018) (4 chains, 1000 warmup and 4000 sampling iterations) using a Bernoulli distribution (logit link) and default uninformed priors. We estimated the Bayes factors with the “bayes_factor” function of the “brms” package. For post-hoc analyses of the performance on the implicit memory test, we computed X^2^ tests. Corresponding Bayes factors were obtained using the “proportionBF” function (R-package “BayesFactor”, v0.9.12-4.2; Morey et al., 2018)

## 3 Results

Added over all 26 participants, a total of 1149 word-pairs was played during SWS. In a subtotal of 304 word-pairs, the second word of a pair hit a slow-wave peak twice, three or four times in the four repetitions of a word-pair. In a subtotal of 845 word-pairs, the second word of a pair never or only once hit a slow-wave peak in the four repetitions of a word-pair.

Participants’ guessing accuracy on the implicit memory test was influenced by the number of times the second word of a sleep-played word-pair hit a slow-wave peak (effect of peaks (0/1 vs. 2-4): *X_W_^2^*(1) = 6.848, *p* = .009, *BF_10_* = 10.474). Accuracy exceeded the chance level of 50% (accuracy: 58.2%, *X^2^*(1) = 8.223, *p_Bonferroni,2 tests_* = .008, *BF_10_* = 8.064), if the second word of a pair had coincided with a slow-wave peak in at least two of the four repetitions (0/1 peaks: accuracy = 49.6%, X^2^(1) = 0.058, *p_Bonferroni,2 tests_* > .999, *BF_10_* = 0.088) (Fig. 2A)(Züst et al., 2019). Hence, repeated peak-associated stimulus presentation provided optimal conditions for sleep-learning.

**Figure 2.**
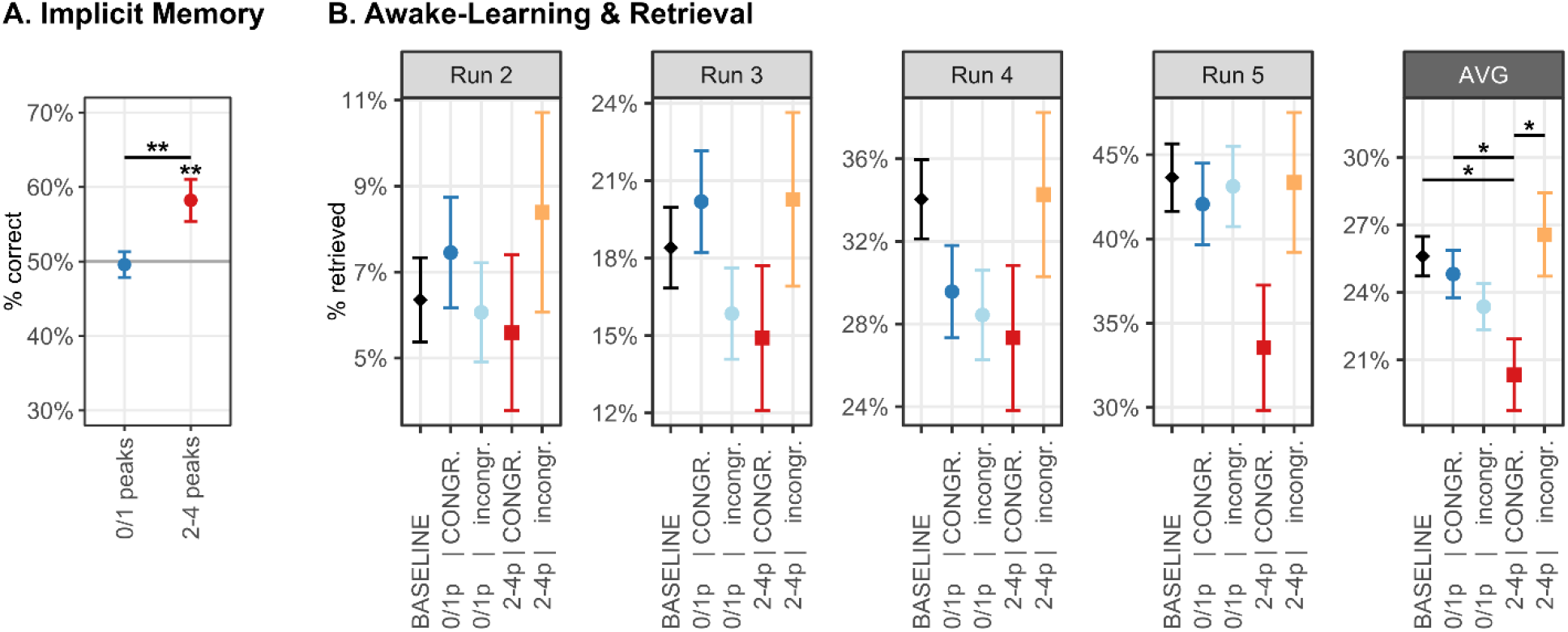
Results. A) Implicit memory performance: % correct responses. Response accuracy exceeded chance performance (50%) for word-pairs whose second word was played during a slow-wave peak more than once (2-4 peaks). B) Retrieval performance in the awake-learning and retrieval task: % correctly retrieved associations. Performance is plotted separately for run2 to run5. The rightmost panel (AVG) indicates the average of retention across the four runs. Conditions: BASELINE = new word-pairs; CONGR. = congruent condition; incongr. = incongruent condition. Retrieval performance is displayed for word-pairs whose second word had been sleep-played more than once (2-4p) during a slow-wave peak (optimal conditions for sleep-learning) and for pairs whose second word had never or once (0/1p) been played during a slow-wave peak (poor conditions for sleep-learning). Plots display % correct responses (A) and % correctly retrieved associations (B) at the single-trial level. Error bars indicate binomial standard errors. **p<.01, *p<.05.

Participants’ recall performance on awake-learning increased from run to run *(X_W_^2^(1)* = 709.919, *p* < .001, *BF_10_* > 10^3^). Condition modulated recall performance (congruent/incongruent/baseline; *X_W_^2^(2)* = 1.004, *p* = .604, *BF_10_* = 0.060) only in interaction with the number of times the second word of a sleep-played pair hit a slow-wave peak (interaction: condition*peaks: *X_W_^2^*(1) = 3.947, *p* = .047, *BF_10_* = 3.235)(Fig. 2B). Wake learning in the congruent condition was paradoxically diminished (rather than enhanced) when conditions for sleep-learning had been optimal (second word repeatedly hit peaks across repetitions) versus poor (second word hit peak once) (*X_W_^2^*(1) = 6.676, *p_Bonferroni, 4 tests_* = .039, *BF_10_* = 10.493). Awake-learning in the congruent versus incongruent condition was diminished if sleep-learning conditions had been optimal (*X_W_^2^*(1) = 6.330, *p_Bonferroni, 4 tests_* = .048, *BF_10_* = 9.556), but not if they had been poor (*X_W_^2^*(1) = 0.132, *p_Bonferroni, 4 tests_* > .999, *BF_10_* = 0.265). However, awake-learning in the incongruent condition was not modulated by the conditions during sleep-learning (peaks: *X_W_^2^* (1) = 0.745, *p_Bonferroni, 4 tests_* > .999, *BF_10_* = 0.515). Moreover, awake-learning in the congruent condition following optimal sleep-learning conditions was also diminished when compared to the awake-learning performance in the baseline condition, where word-pairs had not been played during sleep *(X_W_^2^*(1) = 6.325, *p_Bonferroni, 4 tests_* .048, *BF_10_* = 8.174). The retrieval performance in the congruent condition following optimal sleep-learning conditions dropped more than 10% in the fifth run (33.5% vs. 43.6% recalled associations). Neither recall performance in the incongruent condition nor recall performance in the congruent condition following poor sleep-learning conditions differed from the recall performance in the baseline condition (all *p_Bonferroni, 4 tests_* > .999, all *BF_10_* < 0.360).

## 4 Discussion

We explored whether sleep-learning expedites awake-learning of semantically congruent associations. Paradoxically, sleep-learning diminished rather than promoted awake-learning of semantically congruent associations but left the awake-learning of incongruent associations uninfluenced. Although the encoding of information that is congruent/incongruent to stored information is known to promote/diminish memory formation (Höltje et al., 2019; van Kesteren et al., 2010), the present switchover from the sleeping to the waking state has apparently altered this interaction. The altered interaction is not unprecedented: Andrillon et al. (2017) gave their participants a perceptual learning task during slow-wave sleep and again following awakening. Participants’ awake ability to learn to detect repeating noise patterns in the stream of random noise was impaired if the same noise patterns had been presented during SWS, but not if the patterns were new or had been presented during REM sleep or light non-REM sleep (Andrillon et al., 2017). What was the mechanism that lowered awake-learning following sleep-learning? According to Andrillon et al. (2017), stimulus presentations during SWS were associated with a stronger-than-normal synaptic downscaling in neurons that are activated by the sleep-played stimuli. Synaptic downscaling renews the learning capacity of the brain by eliminating unused and irrelevant synapses (Tononi & Cirelli, 2014). Andrillon et al. (2017) surmised that an exaggerated downscaling may have impaired their participants’ ability to process and store the sleep-played information following waking. Yet, the present data are incompatible with an exaggerated downscaling in stimulus-specific networks because participants had learned the sleep-played vocabulary and retrieved it on the implicit memory test administered following awakening. Learning mostly requires synaptic potentiation (Takeuchi et al., 2014), which is probably why sleep-learning in the present study was successful only during peaks of slow-waves when synaptic potentiation is possible (Andrillon & Kouider, 2020; Destexhe et al., 2007). We therefore assume that semantic associative learning had interfered with the normal synaptic downscaling in those cortico-hippocampal neurons that were activated during sleep-learning and - due to their semantic specialization - again activated during awake-learning of the same/similar semantic associations. Indeed, neurons can be spared from downscaling if they are highly active during sleep (Gulati et al., 2017; Tononi & Cirelli, 2020). If neurons that were dually involved in sleep-learning and awake-learning had escaped downscaling, their learning capacity is deemed to be insufficient for normal awake-learning. The fact that awake-learning of new semantic associations in the incongruent condition was normal further suggests that the escape from downscaling has only then detrimental effects, when neurons are dually recruited for sleep- and awake-learning due to their semantic specialization. If this interpretation is correct, a recovery nap between sleep-learning and awake relearning might restore synaptic homeostasis (Mander et al., 2011) and learning capacity, which might then leave awake-learning unimpaired or enhance it. These predictions regarding synaptic potentiation could be tested in future animal experiments at both the cellular and behavioral level.

## 6 Acknowledgments

This work was supported by the Interfaculty Research Cooperation grant “Decoding Sleep: From Neurons to Health and Mind” (to K.H.) from the University of Bern and the Swiss National Science Foundation (SNSF) grant P0BEP1_148941-1 (to M.A.Z.).

## 7 Author contributions

Conceptualization, M.A.Z., S.R., and K.H.; Methodology, S.R., M.A.Z., and K.H.; Investigation, M.A.Z. and S.R.; Formal Analysis, SR; Resources, S.R., M.A.Z., and K.H.; Writing - Original Draft, S.R.; Writing - Review & Editing, S.R., M.A.Z., and K.H.; Visualization, S.R.; Supervision, K.H.; Project Administration, K.H.; Funding Acquisition, K.H., and M.A.Z.

## 9 Data availability

The data and analyses relevant for this study are available on https://doi.org/10.17605/OSF.IO/ZSG5F

